# Assessment of 3D MINFLUX data for quantitative structural biology in cells

**DOI:** 10.1101/2021.08.10.455294

**Authors:** Kirti Prakash, Alistair P. Curd

## Abstract

MINFLUX is a promising new development in single-molecule localization microscopy, claiming a resolution of 1-3 nm in living and fixed biological specimens. While MINFLUX can achieve very high localisation precision, quantitative analysis of reported results leads us to dispute the resolution claim and question reliability for imaging sub-100-nm structural features, in its current state.

---

In 2017, Balzarotti et al. (2017) introduced MINFLUX to localize individual fluorophores by probing the emitter with a doughnut-shaped excitation beam. The authors attained a precision ∼1-nm and resolved loci on DNA origami placed 6-nm apart. In 2020, Gwosch et al. (2020) extended the method to fixed and living biological cells, and into 3D and two colors. Using nuclear pore complexes (NPCs) as an example, the authors measure localization precisions of 1-3 nm and assert (1) that MINFLUX can clearly resolve the eightfold symmetry of Nup96 in single nuclear pores; (2) Nup96 is distributed along a ring of 107 nm in diameter; and (3) that 3D MINFLUX can resolve the parallel cyto- and nucleoplasmic layers of Nup96 in single pore complexes, ∼50 nm apart in the axial (*z*) direction.

However, we noticed that little quantitative evidence was given for these claims and have therefore ourselves analyzed the datasets provided by the authors. We independently agreed with their main localisation precision results, but found (1) that the eightfold symmetry of NPCs is rarely visible at a single nuclear pore level and was not clearly determined in structure-based modelling of the localisation datasets; (2) that the mean or best-fit Nup96 ring diameter varies between datasets and the spread of diameters in each dataset is broader than that found by dSTORM (Thevathasan et al., 2019); and (3) the average *z*-distance between cyto- and nucleoplasmic layers of Nup96 localisations was 40.5 nm instead of ∼50 nm, in the dataset on which this claim was based. Furthermore, in 2-color imaging, the inner ring found in similar dSTORM experiments at 40-nm diameter (Löschberger et al., 2012; Thevathasan et al., 2019) was not resolved as a ring by MINFLUX. In summary, our analysis shows that while these MINFLUX datasets demonstrate high 3D precision in localizing molecules, they do not yet appear to demonstrate the accuracy of previously published state-of-the-art dSTORM imaging of NPCs (Thevathasan et al., 2019).

## Per-pore analysis

We first assessed the MINFLUX datasets by estimating the diameter of segmented Nup96 complexes (Fig 1a-c). Qualitatively, Nup96 appeared less well sampled, comparing the number and uniformity of clusters per NPC with previous dSTORM data, and there was a large range of numbers of localisations per NPC (Fig 1e). Using circle fits as described in Thevathasan et al. (2019), we found the diameter distributions in the 2D, 3D 1-color and 3D 2-color datasets (Gwosch et al. (2020) Fig. 1a, 3f, 5c) to be 107 ± 10 nm, 108 ± 7 nm and 111 ± 5 nm (mean ± s.d., *N* = 20), respectively. The Nup96 ring diameter in a dataset was therefore not simply 107 nm, as stated by Gwosch et al. (2020) and referenced in dSTORM and EM data (Thevathasan et al., 2019; Von Appen et al., 2015), and it had a previously unreported spread. Our estimates for the spread of diameters (Fig 1d) for MINFLUX data were larger than that for dSTORM data (radius 53.7 ± 2.1 nm, so diameter 107.4 ± 4.2 nm, *N* = 2,536 (Thevathasan et al., 2019)). We conclude that MINFLUX does not yet outperform dSTORM for this type of measurement and scale of biological feature.

**Fig. 1.**
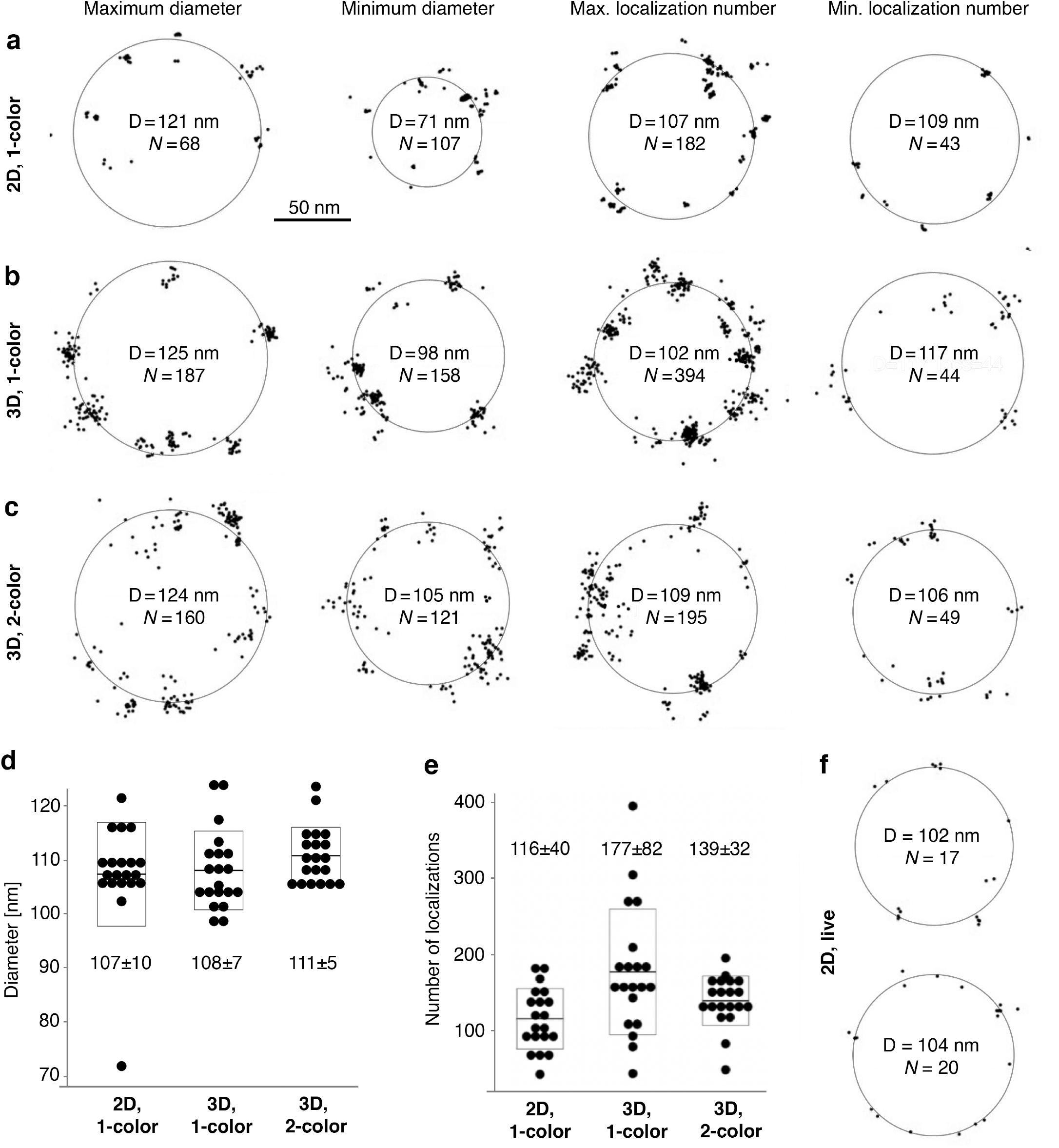
Visualisation of individual nuclear pores. Scatter plots showing localisations from single Nup96 complexes for 2D, 1-color (a); 3D, 1-color (b); 3D, 2-color (c); and 2D, live (f) MINFLUX datasets. We segmented *N* = 20 NPCs for each dataset (a-c) and show those with minimum/maximum diameter and minimum/maximum number of localisations for the outer rings of Nup96. The distributions of the fitted diameter (d) and the number of localizations (e) among the NPCs. Mean ± s.d. is stated and shown. Two Nup96 complexes were visible in the live data (f).

For 2-color, 3D MINFLUX imaging of the NPC, Gwosch et al. show labelled wheat germ agglutinin (WGA-CF680) residing inside the Nup96 octamer both laterally and axially. However, while dSTORM has previously resolved an inner ring of the NPC with WGA-CF680 and measured a diameter of 41 ± 7 nm with WGA-AF647 (Löschberger et al., 2012; Thevathasan et al., 2019), such structure was neither apparent in the MINFLUX data (Fig. S1) nor discussed. Even after segmentation and closer inspection of WGA distributions, we could not visually discern a ring-like structure (Fig. S2). Therefore, we conclude that for this particular sample, MINFLUX, unlike dSTORM, failed to resolve a ∼40 nm ring structure.

## FOV ensemble analysis

We next analyzed the distribution of MINFLUX localizations in a field of view (FOV) using PERPL, a structure-based modelling technique designed for incomplete data, as is often the case for single-molecule localization microscopy (SMLM) (Curd et al., 2020) (Fig. 2). Specifically, we calculated the relative position distribution (RPD) of Nup96 localizations and used its components in the lateral (*xy*) (Fig. 2a–f) and axial (*z*) directions (Fig. 2g–j).

**Fig. 2.**
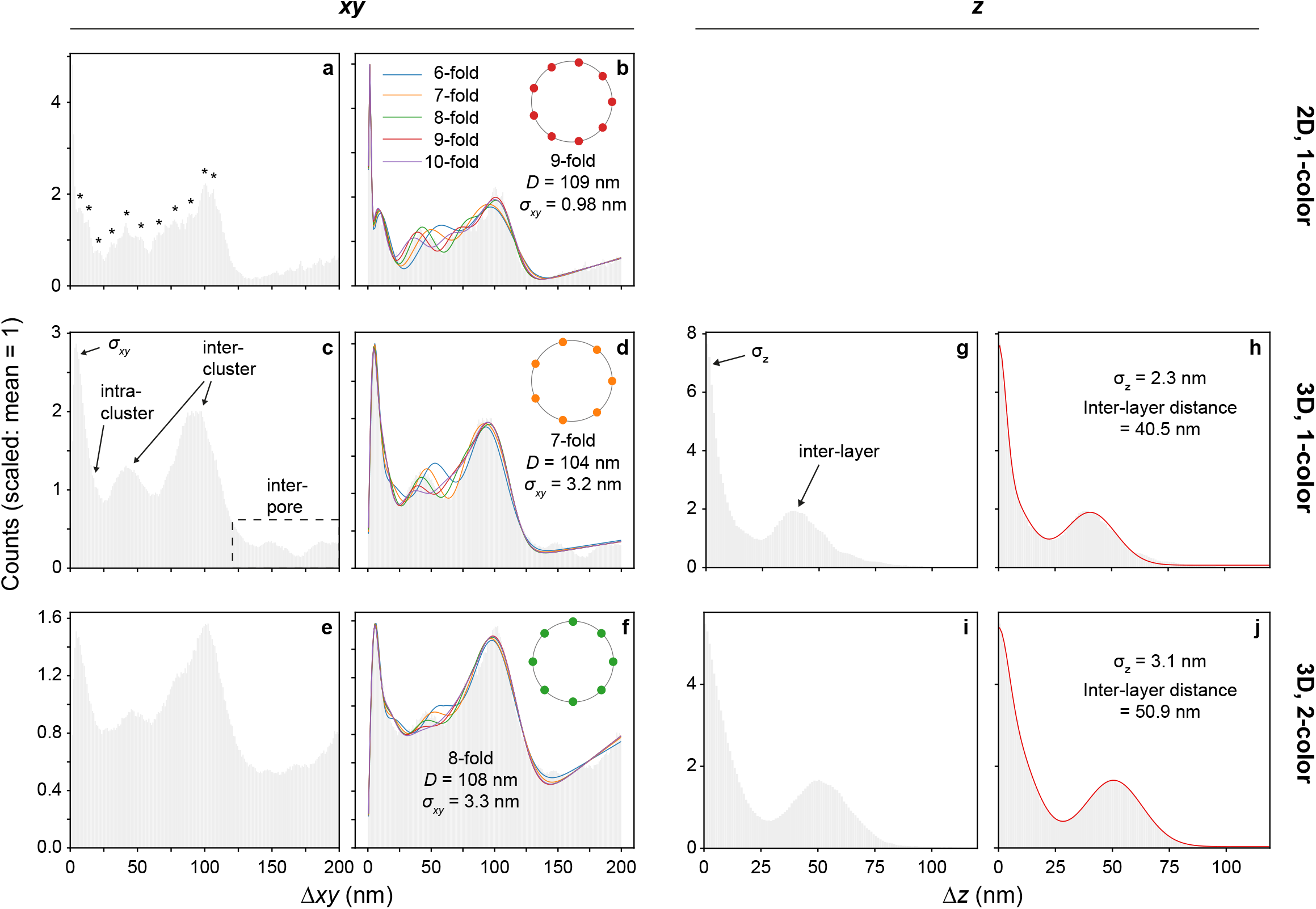
Relative position distributions and model analysis. Histograms of *xy-* and *z-*distances (∆*xy*, ∆*z*) between localisations, bin-width 1 nm. Counts scaled to a mean of 1 to optimise the performance of the fitting algorithm. ∆*xy* distribution for the Nup96 localisations of Gwosch et al. (2020) Fig. 1a (a), 3f (c) and 5c (e) and fits to them (b, d, f,) of nuclear porin models from 6-to 10-fold symmetry, including repeated single-molecule localisations (*σ*_*xy*_), intra- and inter-cluster distances within a NPC, and background≈inter-pore distances (Curd et al., 2020). Symmetry, nuclear pore diameter (*D*) and *σ*_*xy*_ for the model selected by AICc (Curd et al., 2020) in each experiment (b, d, f). Indications of resolved intra-cluster substructure in a (*). ∆*z* distribution for the data in Gwosch et al. (2020) Fig. 3f (g) and 5c (i) and fit with a model including two layers of localisations and repeated single-molecule localisations (*σ*_*z*_) (h, j).

For structures in *xy*, a model RPD that performed well for dSTORM localisations of Nup107 (Curd et al., 2020) was fit to the histogram of the experimental distances between MINFLUX localizations (Fig. 2a–f). Localization precision estimates (*σ*_*xy*_) were 0.98 ± 0.02 nm, 3.20 ± 0.05 nm and 3.31 ± 0.08 nm (fitted value ± 1 s.d. uncertainty) for the 2D, 3D 1-color and 3D 2-color datasets, respectively, of Gwosch et al. (2020), which broadly agree with the published analysis. Secondary peaks may indicate consistent substructure within clusters in the 2D dataset down to a distance of 7 nm (Fig. 2a, not modelled). However, this detail is lost in the 3D and 2-color datasets (Fig. 2c,e). If *σ*_*xy*_ is in the range 1–3 nm, we question whether resolution is possible at 1-3 nm, as is claimed. For instance, FWHM ≈ 2.355*σ*, so *σ*_*xy*_ of 1-3 nm implies FWHM of 2.4-7.1 nm for a single molecule, and we would not expect to clearly resolve molecules closer than this.

Estimates of the Nup96 ring diameters varied from 104 to 109 nm and orders of symmetry from 7-to 9-fold, for the different datasets (Fig. 2a-f). The best fits do not follow the experimental distance distribution as closely as for Nup107 dSTORM data (Curd et al., 2020), which may be due to the effect of intra-cluster substructure (Fig. 2a, b), a more variable arrangement of Nup96, a smaller number of NPCs in the FOV, or the filtering applied to the data. In particular, filtering out a larger fraction of localisations (Fig. 3) appears to have caused the background to deviate from linear (Fig. 2a,b) to a more complex distribution (Fig. 2c–f). These analyses are not consistent with the claim that MINFLUX obtains a diameter of exactly 107 nm and 8-fold symmetry, as reported by Gwosch et al., who did not perform any structural analysis in *xy*. Such conclusive results may be difficult to obtain from these datasets, with small numbers of NPCs (*N* ∼ 20-30, including incomplete complexes), and larger datasets would be useful to establish them.

**Fig. 3.**
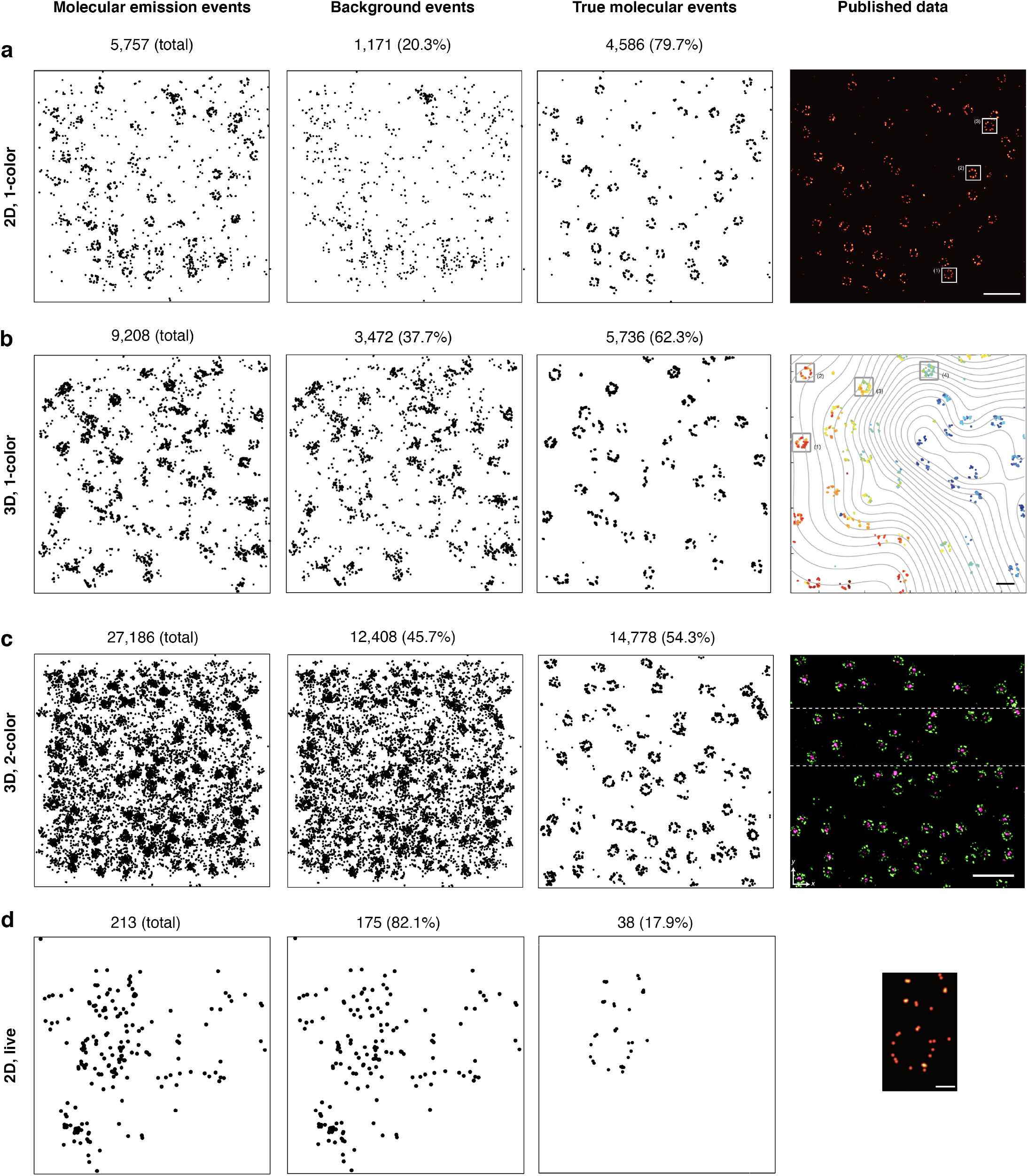
MINFLUX localization filtering. Scatter plots for 2D, 1-color (a); 3D, 1-color (b); 3D, 2-color (c); and 2D, live (d) unfiltered and filtered MINFLUX datasets. The raw MINFLUX data comes in a tabular format, with a Boolean flag indicating that a localisation was assigned as either a background event or a true molecular event. Scale bar: 500 nm (a), 200 nm (b), 500 nm (c), 50 nm (d) in the published data column.

In the axial (*z*) direction, using PERPL (Fig. 2g-j), we estimated localization precision (*σ*_*z*_) at 2.28 ± 0.05 nm and 3.08 ± 0.06 nm for the 3D, 1-color and 2-color datasets of Gwosch et al. (2020), demonstrating the high localisation precision in *z*, and implying that best possible resolution in *z* ≈ 5-7 nm (FWHM). However, for the 3D 1-color dataset of Gwosch et al. (2020) Fig. 3f, we found the distance between the Nup96 layers to be 40.5 ± 0.2 nm (Fig. 2g,h), and not ∼50 nm, as claimed by Gwosch et al. (no quantitative analysis provided) and previously obtained in dSTORM (Thevathasan et al. (2019)), in agreement with EM (Von Appen et al., 2015). In contrast, in the 2-color dataset of Gwosch et al. (2020) Fig. 5c, we estimated the inter-layer distance at 50.89 ± 0.08 nm (Fig. 2i,j), close to the expected ∼50 nm (and estimated at ∼46 nm by Gwosch et al.). Thus, we conclude that MINFLUX does not appear to have reliably provided accurate quantitative measures in the axial direction.

## Live MINFLUX and filtering

The raw MINFLUX data comes in a processed tabular format including the parameters used for event filtering (Sup Table 1). We reproduced the published images of Gwosch et al. (2020) and noted an increased level of filtering from 20% to 83% as we moved from 2D to 3D to live MINFLUX data (Fig. 3). We assume the choice of event filtering parameters in the case of the live data (Fig. 3d) was needed to deal with increased background and movement during localization, and visually appeared to succeed in finding a collection of ≤38 true positive Nup96 localizations. To apply MINFLUX to biological research questions, an explanation of how to select the values of the event filtering parameters is needed. In particular, how to optimize the number and proportion of true positives in an investigation of an unknown sample structure, where this may be challenging.

**Table 1.**
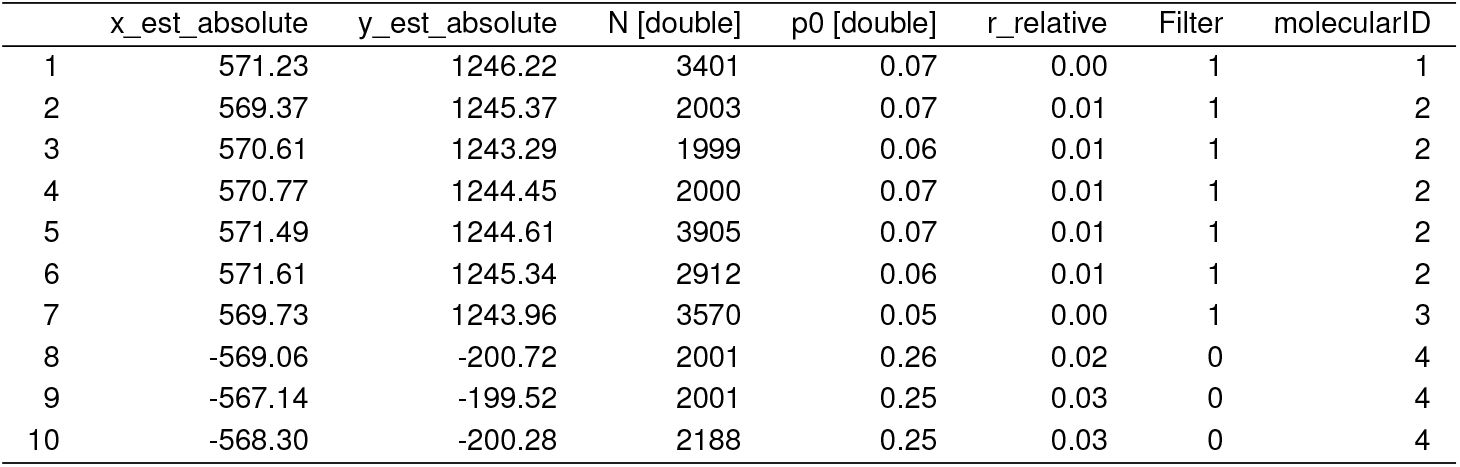
MINFLUX raw datasets in form of localisation positions as provided by the authors (Gwosch et al., 2020)

|  | x_est_absolute | y_est_absolute | N [double] | p0 [double] | r_relative | Filter | molecularID |
| --- | --- | --- | --- | --- | --- | --- | --- |
| 1 | 571.23 | 1246.22 | 3401 | 0.07 | 0.00 | 1 | 1 |
| 2 | 569.37 | 1245.37 | 2003 | 0.07 | 0.01 | 1 | 2 |
| 3 | 570.61 | 1243.29 | 1999 | 0.06 | 0.01 | 1 | 2 |
| 4 | 570.77 | 1244.45 | 2000 | 0.07 | 0.01 | 1 | 2 |
| 5 | 571.49 | 1244.61 | 3905 | 0.07 | 0.01 | 1 | 2 |
| 6 | 571.61 | 1245.34 | 2912 | 0.06 | 0.01 | 1 | 2 |
| 7 | 569.73 | 1243.96 | 3570 | 0.05 | 0.00 | 1 | 3 |
| 8 | -569.06 | -200.72 | 2001 | 0.26 | 0.02 | 0 | 4 |
| 9 | -567.14 | -199.52 | 2001 | 0.25 | 0.03 | 0 | 4 |
| 10 | -568.30 | -200.28 | 2188 | 0.25 | 0.03 | 0 | 4 |

With *N* = 2 NPCs and 38 localizations, there was too little data available to assess the claim of resolution at 1-3 nm in living cells. Circular fits found diameters of 102 nm and 104 nm (Figure 1f, ideal result 107 nm). We are unsure of *σ*_*xy*_ and a related resolution limit in this case since we could not use PERPL analysis on only two instances of the NPC with missing data. Gwosch et al. did not estimate *σ*_*xy*_ and resolution for localizations in the live sample. However, in a fixed sample, they measured *σ*_*xy*_ ∼2 nm for the same label, Nup96-mMaple (Gwosch et al. (2020) Supplementary Fig. 7), which gives a possible resolution limit (≈ FWHM) of ∼5 nm in that case. Therefore, we cannot quantitatively assess resolution and reliability in the case of live samples, although precision and resolution are generally degraded when moving from fixed to live specimens, so we do not expect nanometer (1-3 nm) resolution in the case of live MINFLUX imaging, based on these datasets.

## Discussion

From analysis of the data reported by Gwosch et al., we were unable to confirm that MINFLUX delivers 3D multicolor nanometer resolution (1-3 nm) in fixed and living cells, at the current stage of the technology. In fact, it generated less precise and reliable results than established SMLM methods and appeared unable to resolve a 40-nm ring structure.

3D Nup96-AF647 localisation precision in fixed samples, after event filtering, is impressive at *σ* = 1-3 nm. However, we advise against interpreting localisation precision as resolution, which is intuitively understood as the distance at which two nearby objects can be distinguished, is larger than 2*σ* at a lower limit, and is affected by other factors such as localisation density. Furthermore, these localisation precisions were found after event filtering, which reduces localisation density and may be difficult to perform effectively on an unknown sample. We suggest assessing resolution, detection efficiency and exploration of event filtering parameters on blind samples, to demonstrate the potential of this new technology. For an initial discussion of these issues, see Prakash (2021).

We fully expect MINFLUX methods to continue to improve, as they have done in the powerful iterative and 3D developments already reported (Gwosch et al., 2020). However, we recommend that experimentalists also perform initial testing of the resolution, detection efficiency and exploration of event filtering parameters on potential samples, to the extent this is possible, before choosing MINFLUX over other SMLM techniques.

## Data availability

The MINFLUX data was made available by Stefan Hell. All the re-analyzed data and code will be made available via Zenodo.

## Acknowledgements

We would like to thank Lothar Schermelleh, Johannes Hohlbein and Michelle Peckham for helpful discussions. A.P.C. gratefully acknowledges funding by the UK Biotechnology and Biological Sciences Research Council (BB/S015787/1).

## Supplemental Material

### Methods

In Fig. 1, we used the MATLAB function circlefit (https://uk.mathworks.com/matlabcentral/fileexchange/5557-circle-fit) to fit a circle to a set of (*x, y*) points, All the plots for Fig. 2 were generated using PERPL (https://bitbucket.org/apcurd/perpl-python3/) ((Curd et al., 2020)). The *xy*-model includes Gaussian clusters arranged symmetrically around a ring, repeated localizations of a single molecule (with s.d. spread *σ*_*xy*_) and a linearly increasing background term. The most likely order of symmetry is selected using corrected Akaike information criteria, which are calculated from the residuals of the model fits. The *z*-model includes two layers of localizations, each with a Gaussian distribution in *z*, a term for localization precision (*σ*_*z*_) for repeated localizations of a single molecule, and constant background.

All the re-analyzed data has been deposited to Zenodo.

**Fig. S1.**
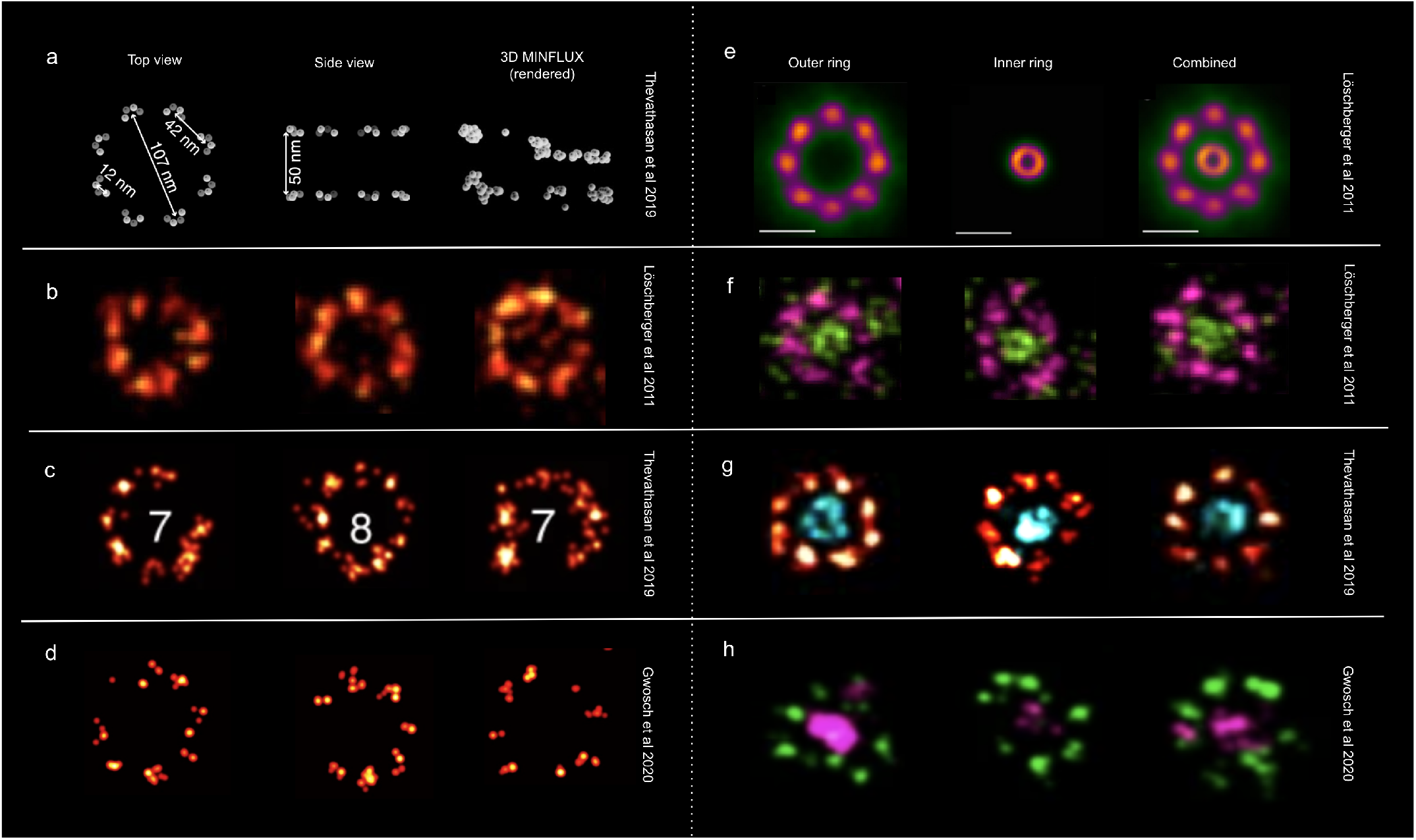
Nuclear pores across different imaging modalities. Figure and caption modified from Prakash (2021). (a) A schematic of the Nup96 complex, taken from Thevathasan et al. (2019). 3D MINFLUX rendered data presented for comparison from Gwosch et al. (2020) (colormap removed for comparison). Note the uneven distribution in *xz*, compared with the EM model (Von Appen et al. (2015)) and dSTORM data in Thevathasan et al. (2019) Fig. 2h. (b) Membrane protein gp210 from amphibian oocytes imaged with dSTORM (Alexa Fluor 647). The 8-fold symmetry and circular structure of NPCs are generally seen. The outer diameter is ∼120 nm and FWHM of gp210 is ∼30 nm. Image adapted from Löschberger et al. (2012). (c) Nup96 endogenously labelled with SNAP-tag-Alexa Fluor 647 in U2OS cell lines. 8- and 7-component pores are more commonly observed. The effective labelling efficiency for SNAP-Alexa Fluor 647 was ∼60%. Image adapted from Thevathasan et al. (2019). (d) MINFLUX imaging of U2OS cell expressing Nup96–SNAP labelled with Alexa Fluor 647. In these cases (selected in Gwosch et al. (2020) Fig. 1a), there are possible indications of 6-8 clusters, but 6- and 7-component nuclear pores are more prominent throughout the FOV, raising a question about detection efficiency. Localisations were rendered with a Gaussian kernal, *σ* = 2 nm, to visualise the 4 individual copies of Nup96 per NPC subunit (1-5 sub-clusters per subunit apparent here). In multi-color MINFLUX imaging, the localizations in a subunit appear as a larger, undefined cluster (**h**). Cell line and labelling strategy as in Thevathasan et al. (2019). Image adapted from Gwosch et al. (2020). (e) Average images of gp210 (outer ring, *N* = 426) and WGA (central channel, *N* = 621) of the NPC. The outer ring (gp210) has an average diameter of ∼120 nm. The diameter of the inner ring (WGA) is ∼40 nm. Image adapted from Löschberger et al. (2012). Scale bar: 100 nm. (f) dSTORM images of WGA labelled with ATTO 520 (green) and gp210 labelled with Alexa Fluor 647 (pink) in amphibian oocytes. Both the outer ring and inner channel are visible (Löschberger et al., 2012). (g) Two-color SMLM image of Nup96-SNAP-Alexa Fluor 647 (red) and WGA-CF680 (cyan) in U2OS cell lines. The outer ring is clearly visible and the inner ring is also visible in most cases. Image adapted from Thevathasan et al. (2019). (h) Two-color MINFLUX imaging of U2OS cell expressing Nup96–SNAP labelled with Alexa Fluor 647 and WGA conjugated to CF680. The subunits of the outer ring, which each have 4 copies of Nup96, now appear as single clusters (comparing with **c**). The inner ring (WGA) also appears as undefined aggregations of signal. Image adapted from Gwosch et al. (2020)

**Fig. S2.**
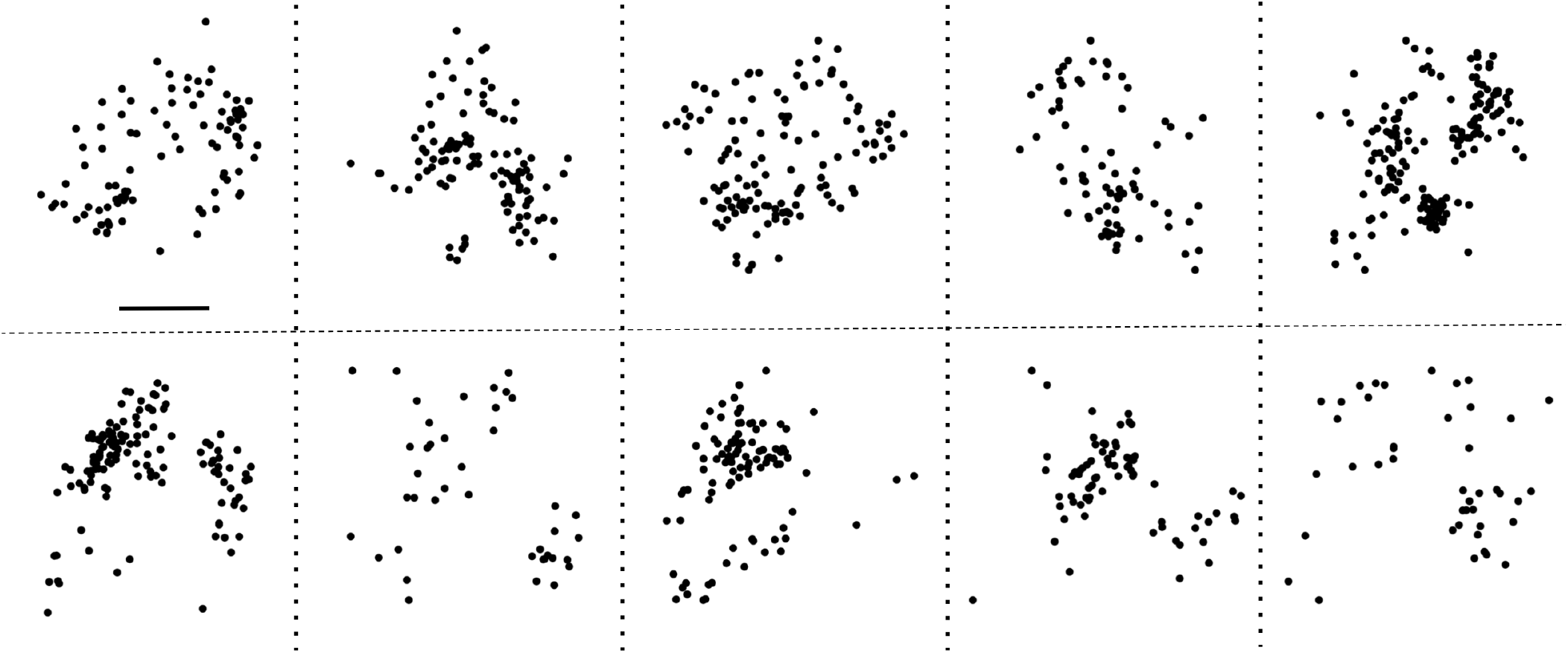
Visualisation of the inner ring of nuclear pores (wheat germ agglutinin (WGA)). Scatter plots showing localisations from 10 segmented WGA complexes from the 3D, 2-color MINFLUX dataset. The segmented complexes did not contain rings of localisations as found by Thevathasan et al. (2019) and Löschberger et al. (2012). Scale bar: 20 nm.

~~~
- x_est_absolute[double]: Absolute estimated molecule position in um.
- y_est_absolute [double]: Absolute estimated molecule position in um.
- N [double]: Number of photons used for localization.
- p0 [double]: Ratio of photons collected for central STC position.
- r_relative [double]: Relative distance of molecule position with respect to the central STC position in um.
- filter [logical] (1-color data only): Boolean flag result of event filters.
         True: localization is valid (molecular emission event)
         False: localization is invalid (background event).
- moleculeID [double]: ID of molecular emission event.
~~~

